# Structural diversity and conservation among CRESS-DNA bacilladnaviruses revealed through cryo-EM and computational modelling

**DOI:** 10.1101/2025.08.04.668413

**Authors:** Johanna Gebhard, Yuji Tomaru, Kenta Okamoto, Anna Munke

## Abstract

Viruses that infect single-celled algae strongly regulate microalgae growth and community composition through cell lysis, enable nutrient recycling in marine ecosystems, and offer valuable insights into early stages of viral evolution. One major group, the *Bacilladnaviridae* family of single-stranded DNA viruses, infects diatoms in marine environments. Here, we present the capsid structure of *Chaetoceros lorenzianus* DNA virus (ClorDNAV, *Protobacilladnavirus chaelor*) determined at 2.2 Å resolution, thereby expanding the known structural diversity within the *Cressdnaviricota* phylum. The ClorDNAV capsid protein (CP) contains a conserved jelly-roll fold and a surface-exposed projection domain, with both N- and C-termini oriented toward the capsid interior. A low-resolution reconstruction of the genome revealed a spooled arrangement of the outer DNA layer, similar to that observed in *Chaetoceros tenuissimus* DNA virus type II (CtenDNAV-II). Structural comparison with CtenDNAV-II revealed five key CP differences: the absence of surface-exposed C-terminal tails in ClorDNAV, the presence of a helical domain, differences in the projection domain conformation, variation in the number of β-strands in the jelly-roll fold, and the lack of ion-attributed densities at subunit interfaces. Together with the genome reconstruction, these findings underscore the importance of experimentally determined structures for understanding viral architecture and evolution. To complement these results, we analyzed AlphaFold3-predicted CPs from all classified *Bacilladnaviridae* genera. These models confirmed the conservation of the jelly-roll fold across the family while revealing variability in the surface-exposed and terminal regions, likely reflecting host-specific adaptations and genome packaging strategies. Together, the experimental and predicted structures provide a comprehensive view of structural conservation and divergence in *Bacilladnaviridae*. Furthermore, the results provide additional structural evidence for the evolution of ssDNA *Bacilladnaviridae* from a noda-like ssRNA virus ancestor and suggest a shared genome organization within *Bacilladnaviridae* that resembles those observed in viruses with double-stranded genomes.

## Background

Viruses effectively regulate microbial communities in marine environments by causing selective mortality of infected hosts. Viral lysis releases carbon and other nutrients as dissolved organic matter, which is subsequently taken up and recycled by other microorganisms (i.e., the ‘viral shunt’ [1–3]). Moreover, many viruses carry auxiliary metabolic genes and act as reservoirs for genetic diversity and as vectors for horizontal gene transfer between microbes [4]. Viral infections thus influence the composition and diversity of microbial populations on a local level, with broader implications for global nutrient cycles [3,5].

Diatoms, a group of unicellular algae, are among the main primary producers in marine environments [6,7]. Their growth is strongly influenced by seasonality, nutrient availability, and predation by both RNA and DNA viruses [8]. In recent years, multiple studies have reported 3D reconstructions of diatom virus capsids using single-particle cryo-electron microscopy (cryo-EM) [9–11]. These models have revealed new insights into how viral capsids bind to the host receptor [9,11], genome packaging, and the evolution of capsid proteins [10].

Here we contribute to this growing body of work by presenting 3D reconstructions of the capsid and outer genome layer of *Chaetoceros lorenzianus* DNA virus (ClorDNAV, *Protobacilladnavirus chaelor*) [12]. ClorDNAV and the previously characterized *Chaetoceros tenuissimus* DNA virus type II (CtenDNAV-II, *P. tenuis*) [10] both have narrow host ranges, infecting only their respective *Chaetoceros* species [12,13]. These viruses are closely related phylogenetically and belong to the *Protobacilladnavirus* genus within the *Bacilladnaviridae* family of single-stranded DNA viruses that infect diatoms. Together with a rapidly expanding number of virus families, bacilladnaviruses are further classified within the phylum *Cressdnaviricota* [15], a group of viruses commonly referred to as circular Rep-encoding single-stranded DNA (CRESS-DNA) viruses.

The genomes of bacilladnaviruses are circular (approx. 3.5-6 kB) and contain short double-stranded regions (<1 kb) of unknown function. These genomes encode at least four open reading frames (ORFs), including the capsid protein (CP) and the characteristic *Cressdnaviricota* replication-associated protein (Rep) [14]. The Rep protein is conserved across *Cressdnaviricota* and contains two functional domains: a His-hydrophobic-His endonuclease and a superfamily 3 helicase, which mediate rolling-circle replication [14–17]. In contrast, the CPs are highly variable [17] and thought to have been acquired from different RNA viruses through multiple independent horizontal gene transfer (HGT) events [15,16,18]. In bacilladnaviruses, the CP is proposed to have originated from an ancestral noda-like virus [16], which is supported by structural evidence from the CtenDNAV-II capsid [10].

The icosahedral capsid of bacilladnaviruses is formed by T=3 asymmetrical units [10], in contrast to the smaller T=1 capsids found in other *Cressdnaviricota* groups [15], presumably reflecting the need to package larger genomes [16]. The bacilladnavirus CP contains the common viral jelly-roll fold that forms the bulk of the capsid and an additional β-sheet projection domain [10,14]. The CtenDNAV-II genome adopts a spooled arrangement inside the capsid [10], possibly involving the partially double-stranded region. While such arrangements are common in non-enveloped icosahedral viruses with dsRNA or dsDNA genomes that use NTP-driven motor proteins for genome packaging [19–21], this was the first such observation for a predominantly single-stranded genome [10]. Notably, bacilladnaviruses with genomes ≥ 4000 nt are enriched in positively charged residues at the N-terminus [14]. These so-called R-arms [22] correlate with genome length and may facilitate genome packaging through interactions with the viral DNA [14].

In this study, we present a 3D reconstruction of the ClorDNAV capsid at 2.2 Å resolution and an atomic model of its CP, thereby expanding the diversity of solved *Cressdnaviricota* capsid structures. We identify shared and divergent features by comparing the ClorDNAV CP to the experimentally determined CtenDNAV-II CP and AlphaFold3-predicted CPs from other *Bacilladnaviridae*. Additionally, we confirm that mature ClorDNAV virions contain a spooled genome arrangement similar to that of CtenDNAV-II, suggesting a conserved mechanism of genome packaging within the family.

## Methods

### Virus production and purification

ClorDNAV was produced as previously described [12]. The algal lysate was further purified by density gradient centrifugation using 15 % to 50 % w/v sucrose gradients, followed by centrifugation at 24,000 rpm (102,170 × *g*) for 18 h at 4 °C (Sw40Ti rotor; Beckman Coulter). All fractions were analyzed by SDS-PAGE, after which selected fractions were pooled and diluted in 13.6 mL of TNE buffer (50 mM Tris pH 7.4, 100 mM NaCl, and 0.1 mM EDTA). The sample was centrifuged again at 28,000 rpm (139,065 × *g*, Sw40Ti rotor; Beckman Coulter) for 2 h at 4 °C. The resulting pellet was resuspended in 100 μL of TNE buffer and concentrated to a final volume of 30 μL using a centrifugal concentrator (5,000 MWCO cutoff, Vivaspin). The final virus concentration was not determined.

### Cryo-EM grid preparation and data collection

A 3.7 μL aliquot of purified ClorDNAV virons was applied to freshly glow-discharged copper grids (Quantifoil R 2/2, 300 mesh, copper), blotted for 3 s at 95 % relative humidity and 4 °C, and plunge-frozen in liquid ethane using a Vitrobot Mark IV (Thermo Fisher Scientific). Images were acquired using a Titan Krios G2 microscope (Thermo Fisher Scientific) operated at 300 kV, equipped with a K3 BioQuantum detector (Gatan) and an energy filter with a 20 eV slit width. Micrographs were collected at a defocus range of –1.2 and –0.4 μm as movies (30 frames, 8 s each, total exposure dose of 45.16 eV Å^−2^) at a magnification of ×105,000, resulting in a raw pixel size of 0.825 Å (see Table S1).

### Image processing and 3D reconstruction

Image processing and single particle analysis were performed with cryoSPARC v4.5.3 [23]. The frames of 9,352 movies were corrected with Patch Motion Correction and Patch Contrast Transfer Function (CTF) Correction. A total of 8,145 particles were selected after template picking and 2D classification. The capsid was reconstructed with imposed icosahedral (I) symmetry after nine rounds of 2D classification. The final map refinement included FSC-mask auto-tightening, minimizing over per-particle scale, negative Ewald Sphere Correction, optimized per-particle defocus, and optimized per-group CTF parameters. The final map reached a resolution of 2.2 Å at the FSC 0.143 threshold [24,25]. The local resolution was calculated in cryoSPARC.

The outer genome layer was reconstructed by following a similar approach to that of Munke et al. 2022 [10]. A mask was generated from the final capsid refinement using cryoSPARC’s Volume Tools Job (threshold 0.45, dilation radius 10 pixels), followed by Particle Subtraction on the final refinement particles. After 2D classification and ab initio reconstruction, a subset of 3,117 particles was selected. The final genome map reached an approximate resolution of 21 Å (at FSC 0.5) and 19 Å (0.143) without imposing symmetry constraints.

### Model building and refinement

The ClorDNAV CP structure was initially predicted from VP2 ([BAJ79015.1](https://www.ncbi.nlm.nih.gov/protein/BAJ79015)) using AlphaFold 2.0 [26] and manually fitted into the cryo-EM density map using Coot [27]. The final model was obtained through iterative rounds of real-space refinement in PHENIX [28] with geometric and secondary structure restraints, combined with manual adjustments in Coot and ISOLDE [29]. Validation statistics are summarized in Table S1. Cryo-EM maps and atomic models were visualized with UCSF ChimeraX v1.9 [30]. The coulombic electrostatic potential of the ClorDNAV and CtenDNAV-II capsids was calculated using ChimeraX. The topology and subunit interactions of the ClorDNAV and CtenDNAV-II capsids were analyzed with PDBsum [31].

### Comparison to other bacilladnavirus capsid proteins

Capsid protein sequences from all 22 classified *Bacilladnaviridae* species were downloaded, translated, and aligned against the ClorDNAV CP as the reference using MUSCLE with default settings [32]. The alignment was visualized using Jalview v2.11 [33], and sequence conservation was analyzed using the ConSurf web server [34,35]. CP structures of the same bacilladnavirus species were predicted using the AlphaFold 3.0 [36] web server with the seed automatically sampled. Representative CPs for each genus were selected for structural comparison, as type species have not yet been designated. All structures were visualized in UCSF ChimeraX v1.9 [30].

### Data availability

The atomic model and cryo-EM map of the ClorDNAV capsid have been deposited in the Protein Data Bank (pdb_00009S3S) and the Electron Microscopy Data Bank (EMD-54550), respectively. The EMDB entry includes the full map, half-maps, and the mask used for refinement. The reconstruction of the outer genome layer has also been deposited in the EMDB (EMD-54548), including the full map, half-maps, and masks for refinement and signal subtraction. Predicted CP structures have been deposited in ModelArchive [37] under DOI: 10.5452/ma-amun-bacp.

## Results

### The ClorDNAV capsid is icosahedral with T = 3 symmetry

The structure of the ClorDNAV capsid was determined using single-particle cryo-EM with imposed icosahedral symmetry (Figures S1A & S1B). The final reconstruction reached a resolution of 2.2 Å (Fig. S1C), with local resolutions ranging from 1.82 to 35 Å (Fig. S1D). A summary of data acquisition, processing, and model validation is provided in Table S1.

The ClorDNAV capsid has T=3 symmetry and consists of 180 CP protomers. The asymmetric unit contains three quasi-equivalent CP subunits, designated A, B, and C (Fig. 1A and 1B), following the nomenclature previously established for CtenDNAV-II [10]. We were able to model residues 46/47 to 388 of the 390-residue CP (Fig. S2). The N-termini are located towards the capsid interior (Fig. 1C) but were not resolved in the density. This unresolved region is enriched in positively charged residues (∼45 %), suggesting the presence of an R-arm commonly found in *Bacilladnaviridae* with larger genomes [14]. The C-termini also point toward the capsid interior (Fig. 1C).

**Figure 1.**
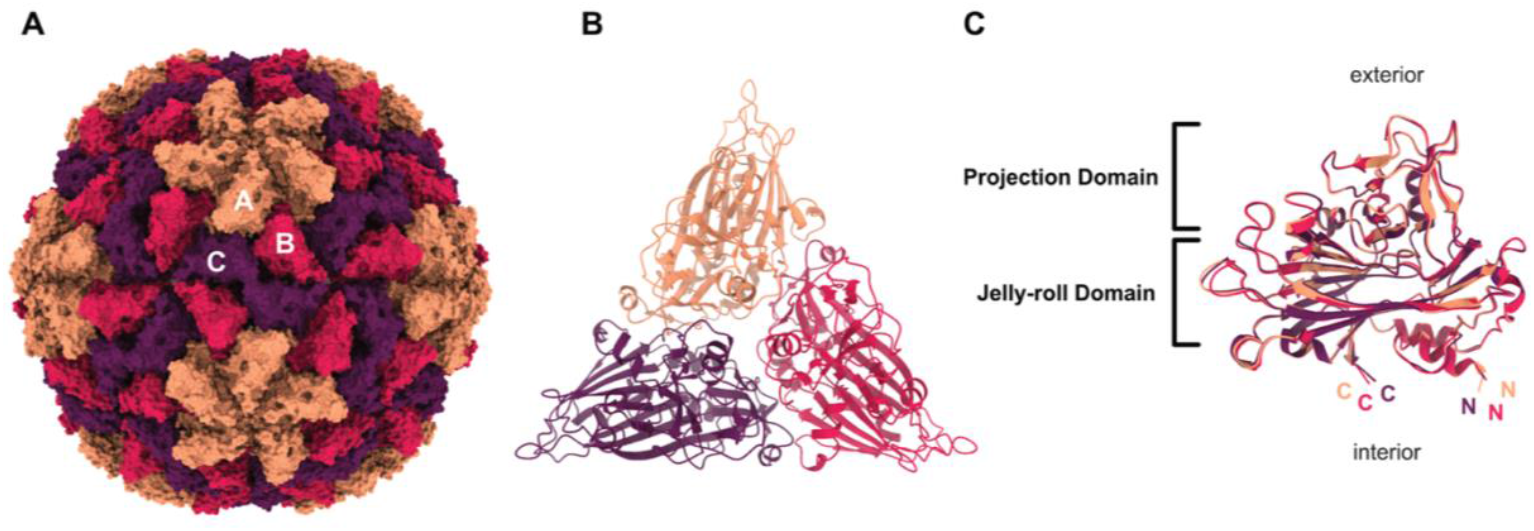
Atomic model of the ClorDNAV capsid. The three subunits A, B, and C of the asymmetric unit are marked orange, red, and purple, respectively. (A) Surface representation of the complete capsid. Secondary structure of a single asymmetric unit viewed from the capsid exterior. (C) Side view of the three superimposed subunits with C- and N-termini labelled.

The three CP subunits are nearly identical (Fig. 1C) and adopt a canonical jelly-roll fold composed of eight β-strands arranged into two antiparallel β-sheets (BIDG and CHEF) [38]. In addition to the canonical strands, the structure includes additional β-strands in each sheet (**I’**BIDG**G’I’’** and **C’’C’**CHEF) (Fig S2). A third β-sheet, composed of seven antiparallel strands formed by loop extensions between strands C-D (2x), E-F (2x), and G-H (3x) (Fig. S2A-B), creates a surface-exposed projection domain (Fig. 1C) [14].

### Atomic model of the ClorDNAV CP reveals structural conservation and divergence within *Bacilladnaviridae*

To assess the structural diversity and conservation within *Bacilladnaviridae*, the ClorDNAV CP was compared to the experimentally determined CtenDNAV-II CP and AlphaFold-predicted CPs from representative species of each *Bacilladnaviridae* genus.

The overall architecture of the ClorDNAV CP is similar to that of CtenDNAV-II [10], with both proteins featuring a conserved jelly-roll fold and a surface-oriented projection domain. However, we identified five key differences between the two viruses’ CPs:

1. **C-terminal orientation:** In CtenDNAV-II, subunits A and B display characteristic 25-residue-long C-terminal tails (Fig. 2A) exposed on the capsid surface (Fig. S3A), whereas in ClorDNAV, all C-termini are oriented toward the capsid interior, similar to subunit C of CtenDNAV-II (Fig. 2A). No additional density was detected in the map that could be attributed to any unmodelled residues. The capsid surface of CtenDNAV-II exhibits a more pronounced coulombic electrostatic charge compared to ClorDNAV, and the C-terminal tails lead to more positively charged grooves on the capsid surface (Fig. S3B). The tails also result in a higher number of predicted inter-subunit interactions in CtenDNAV-II (Fig. S3C).
2. **N-terminal helices:** The N-terminal region is unstructured in CtenDNAV-II subunits A and B but contains α-helices in subunit C and in all ClorDNAV subunits (Fig. 2A). Residues 366 to 377 could not be modelled for CtenDNAV-II subunit C, whereas the equivalent region facing the capsid interior features an α-helix in all three ClorDNAV subunits (Fig. S4). As a result, ClorDNAV contains two interior-facing helices per subunit, whereas CtenDNAV-II contains only one, and only from subunit C.
3. **Projection domain conformation:** Although both viruses possess a projection domain formed by seven β-strands arising from loop extensions between the C-D, E-F, and G-H strands (Fig. S2), the orientations and positions of these loops and secondary structures differ between the two CPs (Fig. 2A).
4. **Jelly-roll fold architecture:** While the overall jelly-roll fold domain between the two viruses is structurally conserved, the number of β-strands differs: the ClorDNAV CP contains five additional β-strands, whereas the CtenDNAV-II CP contains three, relative to the conventional jelly-roll fold [38] (Fig S2).
5. **Subunit interface composition:** In CtenDNAV-II, a density blob at the subunit interface was attributed to a potential metal ion, surrounded by positively charged residues (R84 and R270) (Fig. 2B) [10]. In ClorDNAV, no such density was observed, and positively charged arginine, lysine, or histidine residues within the interface region either face outward (e.g., R80 of ClorDNAV) or are positioned much further apart than in CtenDNAV-II. Instead, residues at corresponding positions are negatively charged (D273) or aliphatic (L72) (Fig. 2C and 2D). Comparison across bacilladnaviruses shows that positively charged residues (K or R) are conserved at position R84 (L72), whereas negatively charged asparagines (D) are conserved at position R270 (D273) instead [14]. Both CtenDNAV-II and ClorDNAV differ from this pattern, with the former having two positively charged residues and the latter neutral and negatively charged residues at the subunit interfaces.

**Figure 2.**
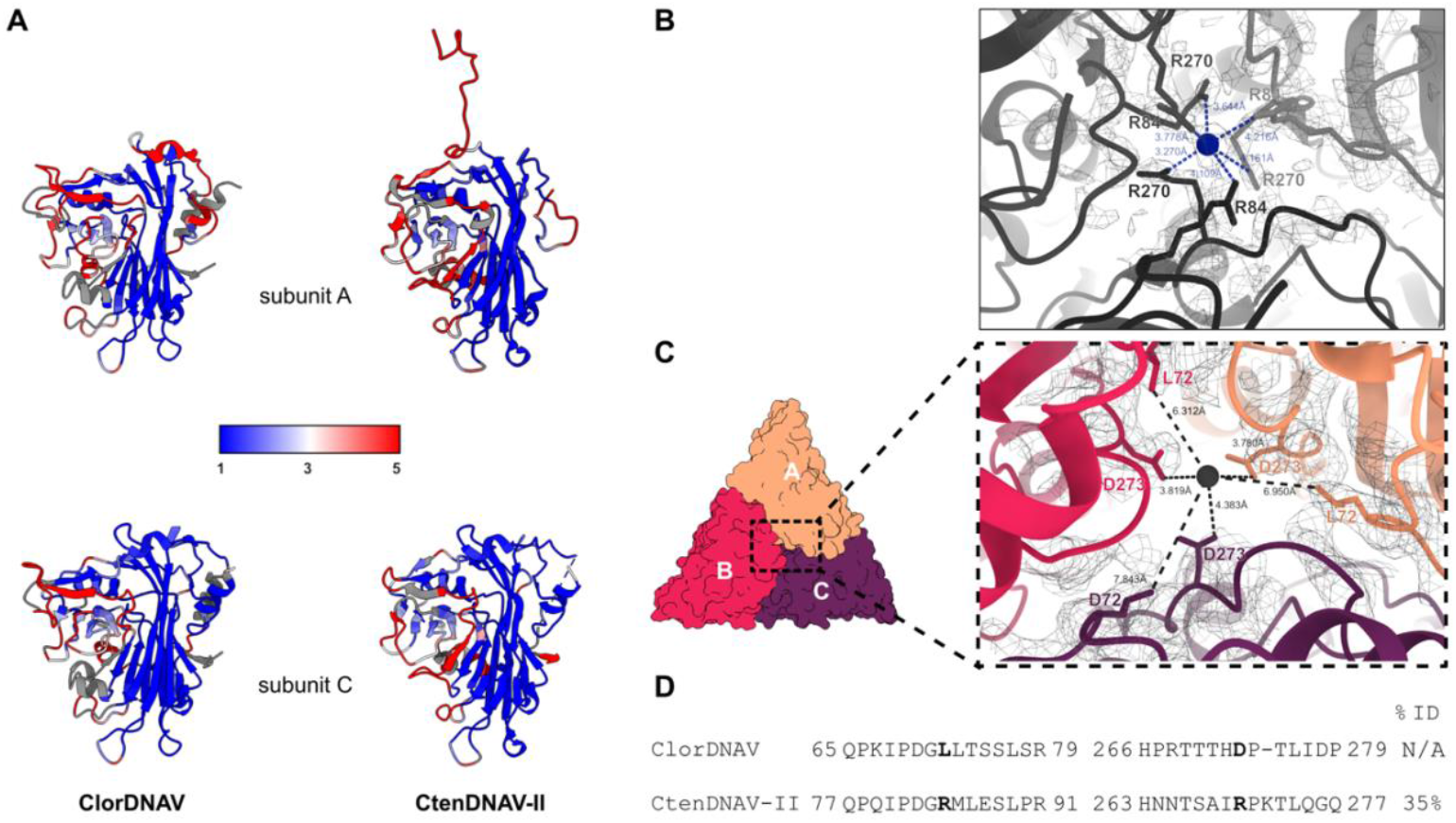
Structural comparison of the ClorDNAV and CtenDNAV-II capsid proteins. (A) Root mean square deviation (RMSD) between ClorDNAV (left) and CtenDNAV-II (right) for subunits A (top) and C (bottom), colored from low (blue) to high (red) deviation; missing values are shown in grey. (B) Subunit interface at the center of the CtenDNAV-II trimer, showing positively charged residues R84 and R270 surrounding unassigned density. (C) Equivalent interface in ClorDNAV, where corresponding residues are either further apart (L72) or negatively charged (D273). Markers were placed in the center of the CtenDNAV-II (blue) and ClorDNAV (black) trimer interfaces to measure the distances (in Ångström) to the surrounding residues with ChimeraX. (D) Sequence alignment of ClorDNAV and CtenDNAV-II CPs, showing selected regions surrounding R84 and R270.

To broaden our structural comparison beyond experimentally determined CP structures, we next examined the AlphaFold3-predicted structures of representative CPs from each *Bacilladnaviridae* genus, based on the updated taxonomy proposed by Varsani et al. [14]. Structural comparisons (Fig. 3A) and analysis of amino acid sequence conservation (Fig. 3B) confirm the pattern observed when comparing the two experimental structures: a highly conserved jelly-roll fold and a more variable third β-sheet which may also be surface-exposed. The C- and N-termini are also more variable and often contain large unstructured regions in the predicted models; we were also unable to model these regions in the CtenDNAV-II and ClorDNAV capsids.

**Figure 3:**
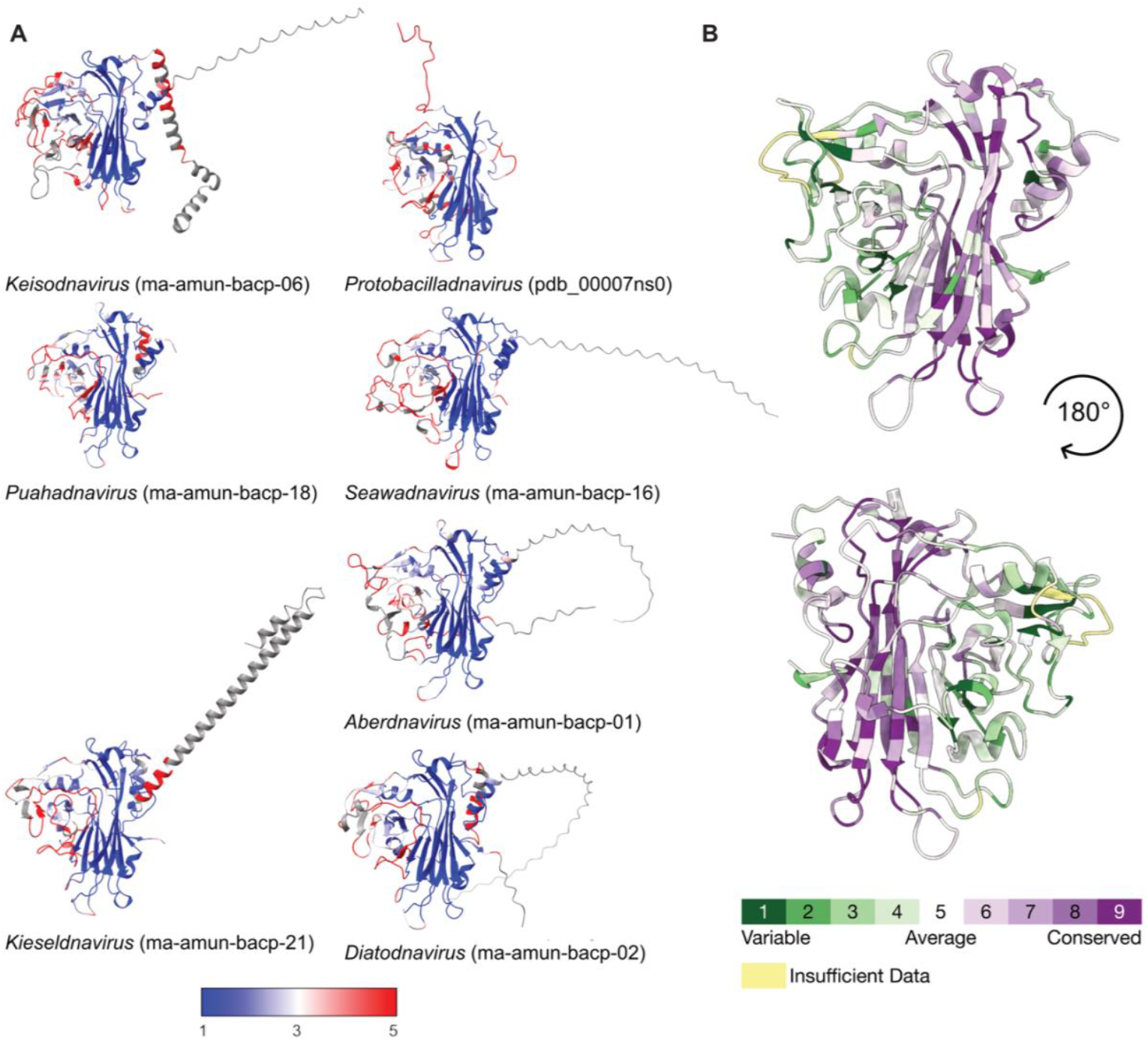
Structure and sequence comparison of bacilladnavirus capsid proteins. (A) Alphafold3-predicted structures of the CPs from one representative species of each *Bacilladnaviridae* genus, according to the latest taxonomy [14]. Structures are colored by RMSD to ClorDNAV subunit A, from low (blue) to high (red) distance; with missing values highlighted in grey. The ModelArchive [37] ID or PDB ID (for CtenDNAV-II [10]) is indicated for each structure. (B) Per-residue conservation scores calculated with ConSurf and projected onto the ClorDNAV CP structure, as indicated by the color scale.

Notably, Avonheates virus SG_120 (*Kieseldnavirus katao*, ma-amun-bacp-21) and *Chaetoceros tenuissimus* DNA virus SS12-43V (*Keisodnavirus chaetenu*, ma-amun-bacp-06) exhibit long α-helices in their respective C- and N-termini (Fig. 3A). Although the terminal ends are modelled with low pLDDT confidence [39], ≤ 50 for AvVSG_120 and ≤ 70 for CtenDNASS12-43V, these modelled extensions could indicate very different capsid organizations compared to the two experimentally resolved *Bacilladnaviridae* CP structures.

### The ClorDNAV genome adopts a spooled arrangement similar to CtenDNAV-II

Previous work has shown that the single-stranded genome of the bacilladnavirus CtenDNAV-II adopts a partially spooled arrangement within the mature virus capsid, although the mechanism of genome packaging remains unknown [10]. In ClorDNAV, higher-order internal structures were already apparent during 2D classification (Fig. S5A-B). The outer genome layer was reconstructed without imposing symmetry (C1) to an approximate resolution of approx. 20 Å (Fig. S5C). Due to the limited number of particles, reconstruction of the internal core was not attempted, although additional densities were visible beneath the outer genome layer. Data processing parameters are summarized in Table S1.

The outer genome layer of ClorDNAV is arranged in a nearly identical spooled configuration to CtenDNAV-II, with turns in the central coil separated by 28 Å (Fig. 4). Although the resolution was insufficient for modelling of the genome arrangement, we hypothesize that, as in CtenDNAV-II, this layer could be formed by the 979 bp partially double-stranded DNA region of the ClorDNAV genome. The spooling pattern appears to cover the entire outer genome layer (Fig. 4, right), but due to the low resolution, we could not confidently assess differences in patterning compared to CtenDNAV-II. Similarly, we were not able to resolve contacts between the capsid and the genome or determine if any non-CPs are partially embedded in this layer.

**Figure 4.**
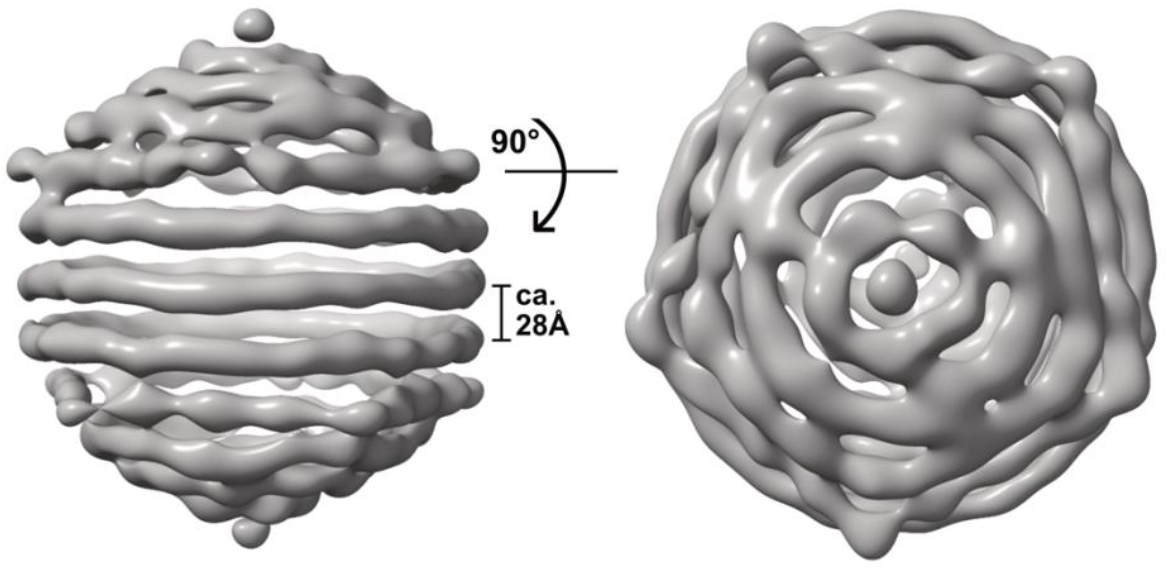
Outer genome layer of ClorDNAV shown in two views, rotated by 90**°**. In the central coil, the parallel turns are separated by approx. 28 Ångström.

## Discussion

The structure of the ClorDNAV virion presented in this study confirms and extends previous findings from CtenDNAV-II [10], demonstrating that key architectural and genomic features, such as the jelly-roll fold and the spooled genome arrangement, are shared across *Bacilladnaviridae*. These features may represent widespread, possibly ancestral traits within the family. At the same time, our comprehensive analysis revealed several structural differences that likely reflect functional divergence or host-specific adaptations.

The ClorDNAV CP consists of a conserved jelly-roll fold formed by two β-sheets, along with a third β-sheet that forms a surface-exposed projection domain. While the jelly-roll is highly conserved across *Bacilladnaviridae*, the projection domain exhibits greater variability, likely due to adaptations to host-specific interactions. In the initial description of the CtenDNAV-II capsid, it was speculated that the CP had been acquired from a noda-like virus ancestor. Specifically, the C-subunit was proposed to resemble the primordial fold, as both termini are oriented toward the capsid interior, similar to what is observed in nodaviruses [10]. Notably, this arrangement is also found in all ClorDNAV subunits, further supporting the idea that this interior-facing orientation may reflect an ancestral structural state within *Bacilladnaviridae*.

Building on the observed differences in terminal orientation, a key distinction between ClorDNAV and CtenDNAV-II lies in the presence of long C-terminal tails in the A and B subunits of the latter. These tails are exposed on the capsid surface, where they contribute to increased inter-subunit interactions and positively charged grooves on the otherwise comparably negatively charged surface. In contrast, ClorDNAV lacks these features; all termini are oriented inward, suggesting that the long C-terminal tails are unique to CtenDNAV-II. However, additional reconstructions of bacilladnavirus capsids are needed to confirm this hypothesis. Sequence comparison between the two viruses did not reveal clear causes for this divergence, implying that the C-terminal modifications in CtenDNAV-II may be linked to host recognition or capsid stability.

In contrast, the N-termini of all subunits in both viruses point toward the capsid interior. The first 46/47 and 59/63 residues of the ClorDNAV and CtenDNAV-II N-termini [10] could not be modelled. These regions are enriched in lysine and arginine residues, suggesting they may function as R-arms that interact with the viral genome [22]. Similar terminal extensions have been shown to mediate genome recognition and packaging in other viruses [22,40,41]. R-arms are common in eukaryotic viruses with icosahedral capsids and ssDNA or ssRNA genomes [42–45]. Interestingly, Varsani et al [14] identified a strong correlation between *Bacilladnaviridae* genome size and the length of the R-arm. It is possible that these R-arms are degraded after genome packaging or that they are bound to the genome in a highly flexible and electrostatically driven manner, which prevents their visualization in the cryo-EM reconstruction. Future higher-resolution reconstructions of the genome layers in CtenDNAV-II and ClorDNAV may resolve the contact points between the capsid and genome. Additionally, structures of bacilladnaviruses with smaller genomes that lack or contain short R-arms [14] could provide important insights into packaging strategies within the family.

The N- and C-termini of the ClorDNAV CP form short α-helices that are positioned parallel to the capsid interior (Fig. S4A). Similar helical domains have been observed in tetraviruses and nodaviruses, where they interact with the viral RNA genomes [46]. Their position in the ClorDNAV capsid is identical to that in previously solved *Alphanodavirus* structures, Fig. S4B [46–48], suggesting a potential role in genome interaction, either in addition to or as an alternative to the R-arm. Although the genome sizes of ClorDNAV and CtenDNAV-II are nearly identical (5,813 nt and 5,770 nt), ClorDNAV contains a longer double-stranded region (979 nt vs. 669 nt). [12,13]. If the spooled outer layer corresponds to this region, ClorDNAV may require stronger interactions between the capsid and the genome, perhaps facilitated by the internally located N and C-termini. However, the current resolution did not allow us to resolve the composition of the outer genome layer or identify possible interactions with the capsid. Nor could we determine whether structural differences exist between the genome arrangements of CtenDNAV-II and ClorDNAV. Nevertheless, the genomic arrangement of the ClorDNAV virion suggests a shared mechanism of genome packaging among ssDNA bacilladnaviruses. This type of organization had previously only been described for dsDNA and some dsRNA viruses [19,20,50], where it is typically generated by NTP-driven motor proteins that actively package the genome into preassembled capsids. However, no homologues of such packaging motors have been identified in bacilladnaviruses to date, raising questions about how these viruses can achieve a similar genome architecture. In parvoviruses, genome packaging is proposed to occur via the helicase domain of a Rep protein, which may transiently associate with the capsid and translocate the genome through the 5-fold pore [51–53]. Bacilladnaviruses may package their genomes through a similar mechanism involving the helicase domain of their *Cressdnaviricota* Rep protein [14–17].

No unmodelled densities were identified in the center of ClorDNAV CP trimers that could be attributed to putative metal ions, known to stabilize capsids in other viruses [54–56]. Combined with the absence of the C-tails, this suggests that the ClorDNAV capsid may be less stable than that of CtenDNAV-II. However, the initial experimental evidence suggested that the ClorDNAV CP trimer has a greater physical stability compared to other ssDNA diatom viruses [12], necessitating further experimental validation. Interestingly, CtenDNAV-II and ClorDNAV seem to be atypical for bacilladnaviruses in different ways. Neither shows the typical pattern of a positively charged residue at position R84/L72 and a negatively charged residue at position R270/D273. Further structural characterization of diverse *Bacilladnaviridae* capsids will be needed to determine larger patterns in the subunit interface organization of bacilladnaviruses. The *Keisodnavirus* and *Kieseldnavirus* genera are particularly interesting, as their respective N- or C-termini are predicted to contain large α-helical extensions, which could have a major impact on the capsid organization.

## Conclusions

Using single-particle cryo-EM, we reconstructed the ClorDNAV capsid and outer genome layer, thereby expanding the known structural diversity of the *Bacilladnaviridae* family. Comparison with the previously solved CtenDNAV-II structure [10] revealed five key differences (Fig. 2A): (1) the positions of terminal α-helices facing the capsid interior, (2) the conformation of the projection domain, (3) the number of β-strands in the jelly-roll domain, (4) the orientation of the C-termini, and (5) the absence of ion-attributed densities at CP interfaces. The last two differences could not have been identified from predicted models alone, emphasizing the importance of experimentally determined structures for understanding viral architecture and evolution. AlphaFold3 models predicted conserved jelly-roll folds across all *Bacilladnaviridae* CPs analyzed in this study (Fig. 3A). However, the observed structural variations in the projection domains and N-terminal regions suggest a degree of plasticity that may reflect host-specific adaptations and genome-packaging mechanisms, which remain to be discovered and experimentally verified.

## Author Contributions

Conceptualization, J.G. and A.M.; sample preparation, Y.T.; methodology, J.G., K.O. and A.M.; validation, J.G.; formal analysis, J.G. and A.M.; investigation, J.G., Y.T., K.O. and A.M.; resources, K.O. and A.M.; writing—original draft preparation, J.G., Y.T., K.O. and A.M.; writing—review and editing, J.G. and A.M.; visualization, J.G. and A.M.; supervision, A.M.; project administration, A.M.; funding acquisition, K.O. and A.M. All authors have read and agreed to the published version of the manuscript.

## Acknowledgements

The data were collected at the Cryo-EM Swedish National Facility in Solna, funded by the Knut and Alice Wallenberg, Erling Persson Family, and Kempe Foundations, SciLifeLab, Stockholm University, and Umeå University. We thank Dustin Morado for his help with data collection. We acknowledge the use of the Cryo-EM Uppsala facility for grid preparation and screening, funded by the Department of Cell and Molecular Biology, the Disciplinary Domains of Science and Technology and of Medicine and Pharmacy at Uppsala University.

