## Supplementary material for "Structural diversity and conservation among CRESS-DNA bacilladnaviruses revealed through cryo-EM and computational modelling"

Content:

Figure S1-S5

Table S1

### Supplementary Figures

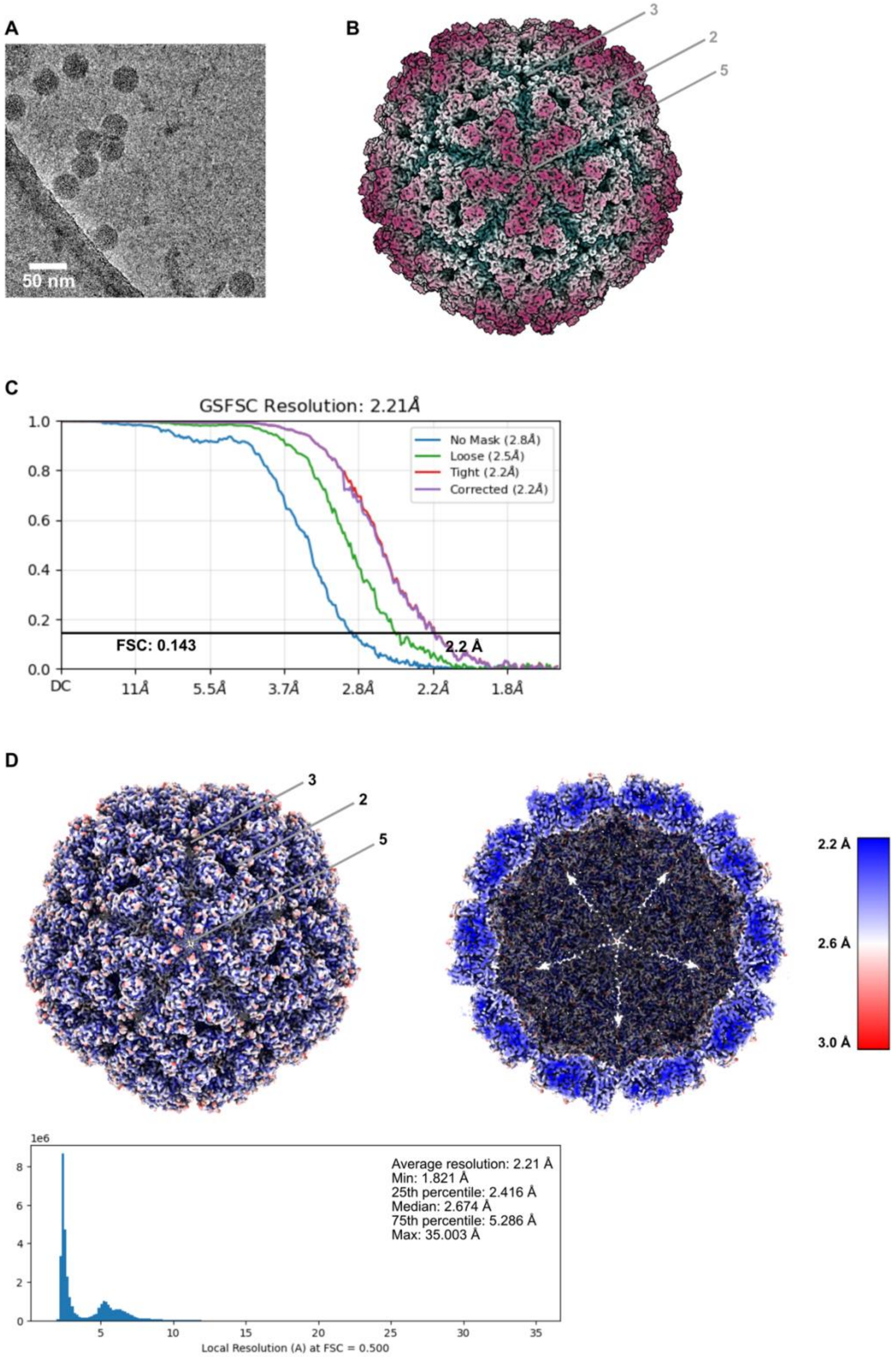

**Figure S1:** Data collection and reconstruction of the ClorDNAV capsid. (A) Raw micrograph of ClorDNAV virions collected during cryo-EM imaging. (B) 3D reconstruction of the ClorDNAV capsid viewed down the 5-fold axis. The icosahedral 5-fold, 3-fold, and 2-fold symmetry axes are indicated as

5, 3, and 2, respectively. (C) Gold standard FSC resolution curves generated by cryoSPARC, showing masked (blue), unmasked (green), FSC-mask tightened (red), and final corrected (purple) curves. The resolution was determined to be 2.2 Å, corresponding to the spatial frequency at which the FSC curve drops below the 0.143 threshold [24,25]. (D) Local resolution map of the final capsid reconstruction, shown from the outside (left) and with the front half removed to show the interior (right). Color scale ranges from blue (2.2 Å) to white (2.6 Å) to red (3.0 Å). Histogram created by cryoSPARC shows voxel distribution by local resolution at FSC = 0.5.

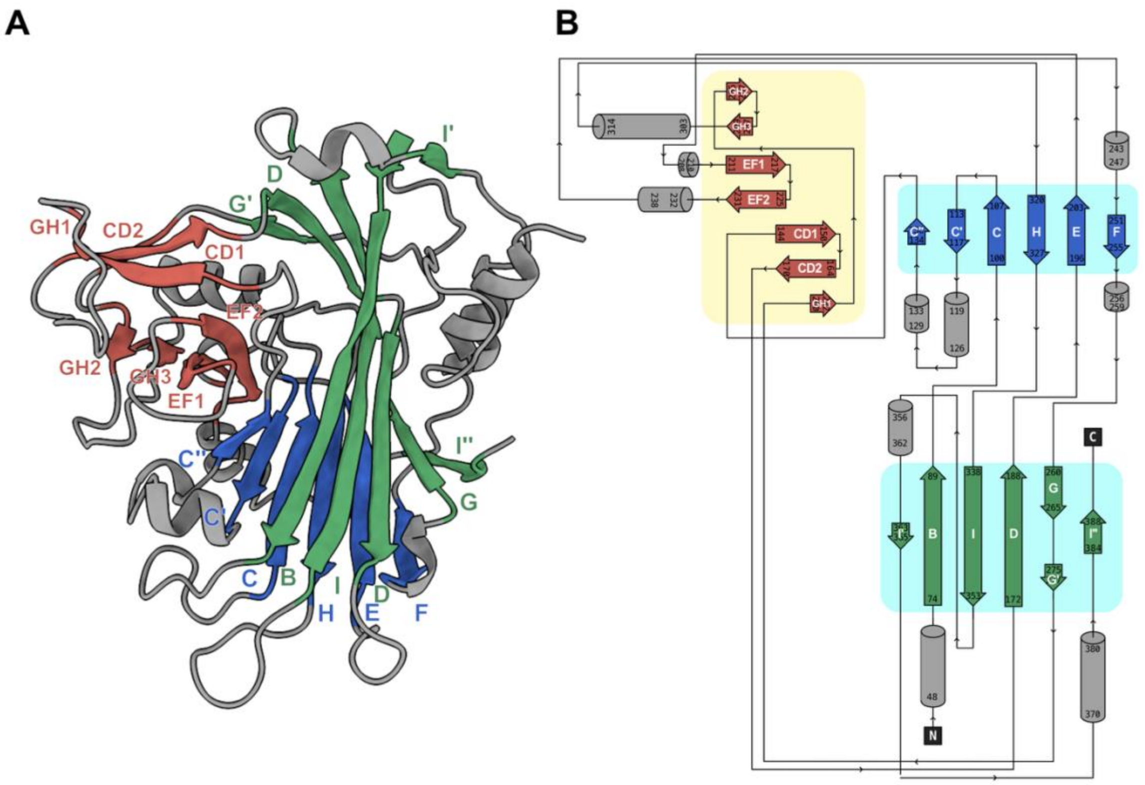

C

MAGRRKQRRS RAPRRKRTTG RKTLKRKADS RATPAKSTRK RTKTVSGAAK HHAVRVKPFS NATTQPKIPD GLLTSSLSRR
LQNVVGVRNG NSPSVHAGSD VMHVVIAPTL GVPVMIANSA EGVLKRPGLS QESSFIGFPG QTVGFENLIE STGVPTWPPT
IPTGQKLENK GGFVLWRIIS QGLRIDLANS DEENDGWFEA CRFNWRNVPR DVCMTPLDGS TTTNSIGIAP NPLWLEEVGY
GMAMVEQPGY KSGLLKDIKK AEFMLHPRTT THDPTLIDPF EYGGSMTSSG GIDNVYYPSD NVSGNAVRFR DMGVDQNMDW
IYIRLHCRPN NGTSSLGSNF LFNVIQNVEV AFNPSSDFAA FQTINKADTK TKMVADGLNN NPDVFNGRRK

**Figure S2:** Detailed structural topology of the ClorDNAV capsid protein. The β-strands are colored according to β-sheets, and named alphabetically (B to I) according to the conventional jelly-roll fold nomenclature in the green and blue β-sheets. (A) Secondary structure of ClorDNAV subunit A. (B)

Schematic diagram showing the jelly-roll domain (light blue) and the projection domain (light yellow). (C) Amino acid sequence of subunit A, starting from residue 1 and divided into blocks of 10 residues. The respective β-strand is marked above the sequence. The first 45 residues, which could not be modelled, are shown in grey and are enriched in positively charged residues (21 total).

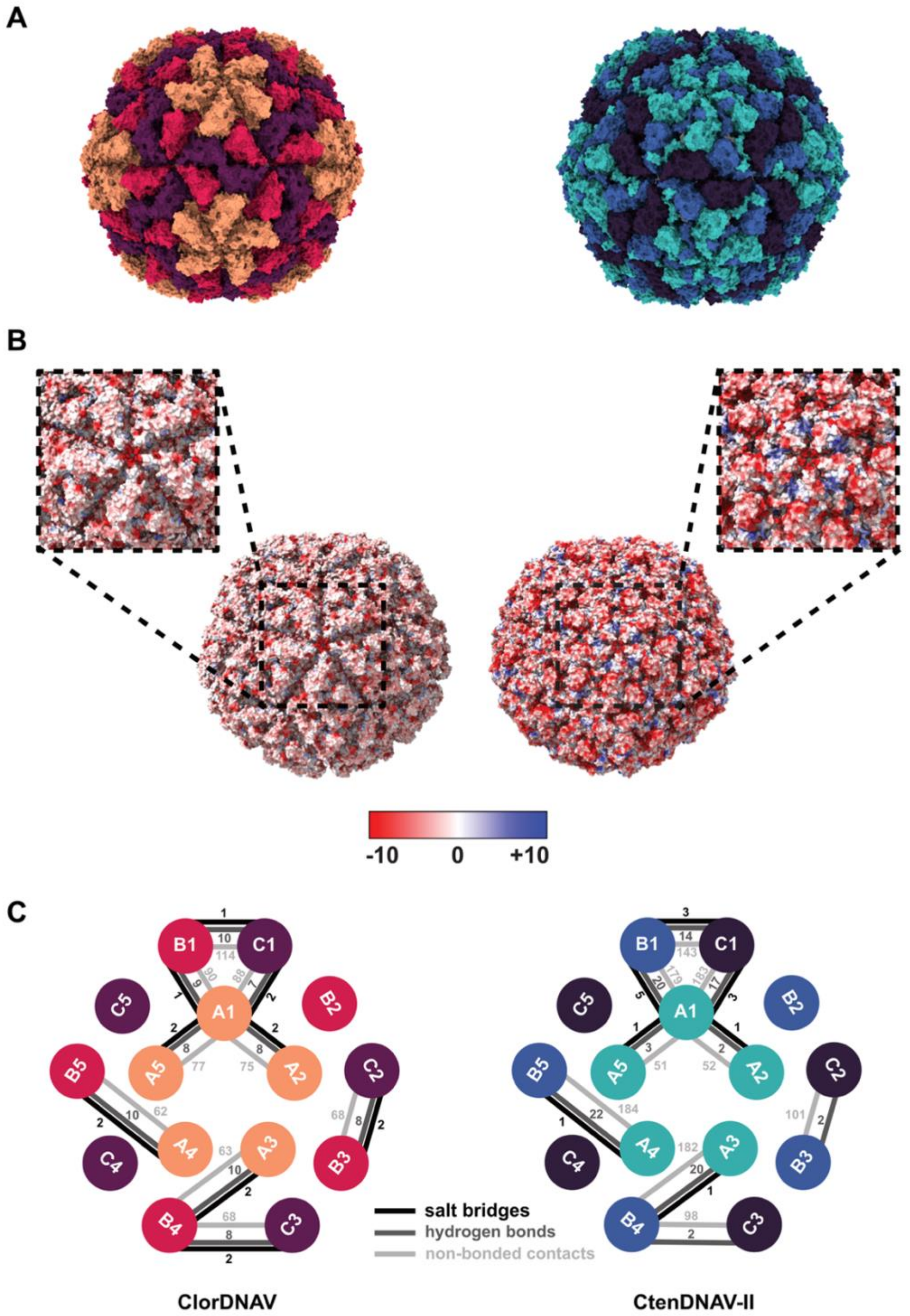

**Figure S3:** Biochemical characterization of the ClorDNAV (left) and CtenDNAV-II (right) capsid [10]. (A) Capsid models colored by subunit. (B) Capsid models colored according to their coulombic electrostatic potential, from red (negative) to white (neutral) to blue (positive). Insets show close-ups of the 5-fold symmetry axis. Analysis was performed with ChimeraX v1.9 [30]. (C) Comparison of subunit interactions based on PDBsum analysis [31], showing the number of predicted salt bridges (black), hydrogen bonds (dark grey), and non-bonded contacts (light grey).

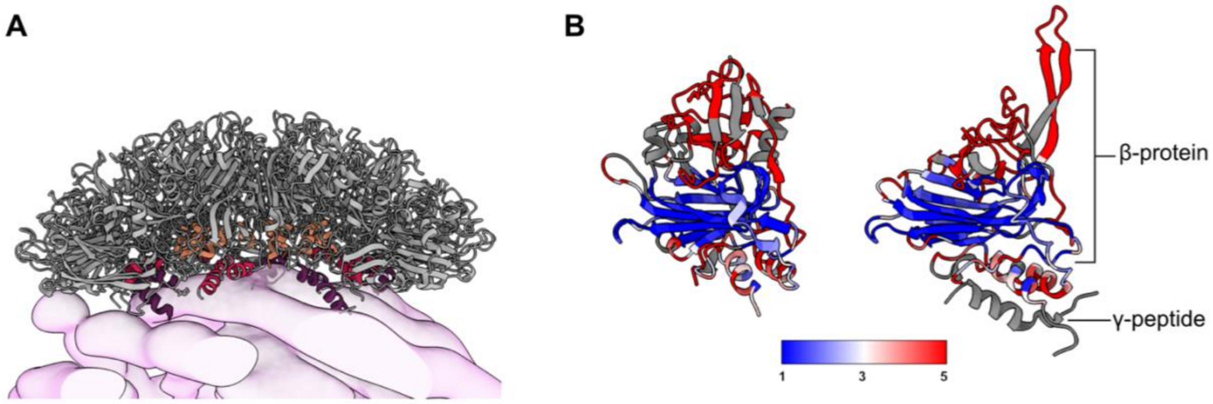

**Figure S4:** The N and C-terminal $\alpha$-helices in the ClorDNAV CP are orientated towards the capsid interior (A) Terminal $\alpha$-helices shown for a ClorDNAV 15-mer, colored by subunit (A: orange, B: red, C: purple). Helices are positioned towards the capsid interior and may interact with the outer genome layer (light pink). (B) Comparison of ClorDNAV subunit A (left) to a Nodamura Virus CP subunit (PDB: 1NOV, right), colored by RMSD from low (blue) to high (red) distance; missing regions are shown in grey. The 1NOV subunit is composed of the β-protein and the γ-peptide [49].

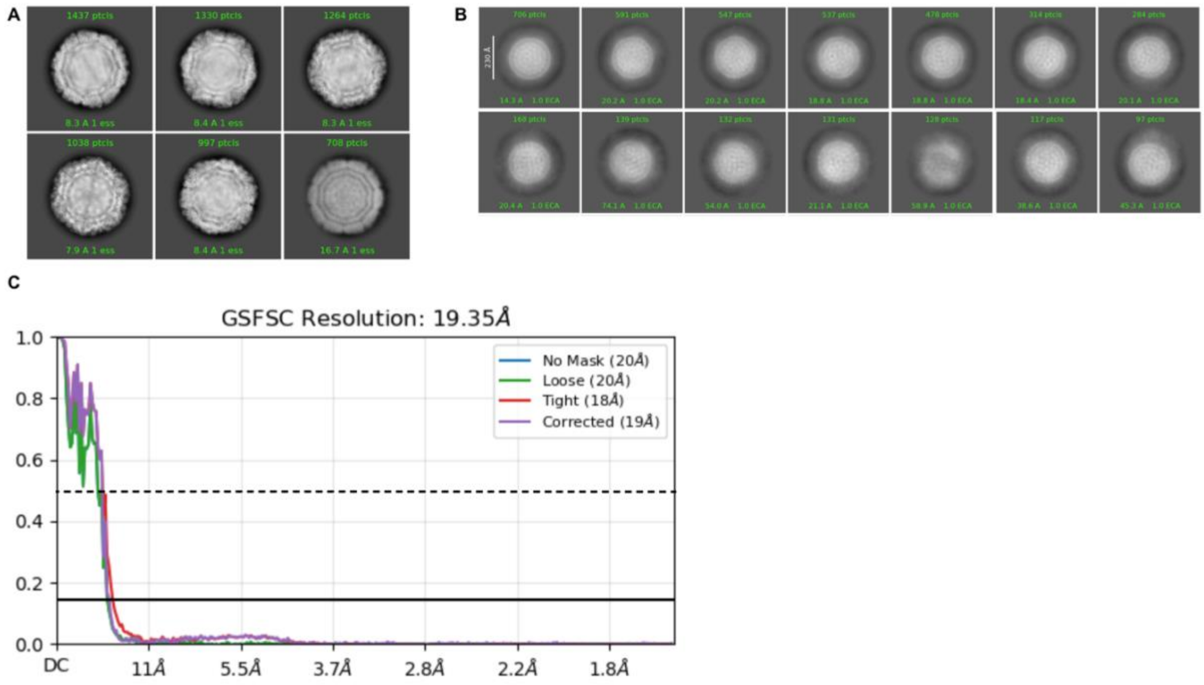

**Figure S5:** Data processing of the outer genome layer of ClorDNAV. (A) Representative 2D classes of ClorDNAV particles showing internal structural features. (B) 2D classes after signal subtraction of the capsid, highlighting genome-associated density. (C) FSC curves of the outer genome layer reconstruction, with estimated resolutions of ~21 Å at FSC = 0.5 (dotted line) and ~19 Å at FSC = 0.143 (solid line).

### Supplementary Tables

**Table S1:** Summary of data collection and processing parameters, along with validation statistics for the model, the data, and model-to-data agreement.

| | ClorDNAV capsid | Outer genome layer |
|---|---|---|
| **Data collection and processing** | | |
| Magnification | 105,000 | |
| Voltage (kV) | 300 | |
| Defocus range (μm) | -0.4 to -1.2 | |
| Microscope | Titan Krios G2 | |
| Camera | K3 BioQuantum | |
| Total electron dose ($e^-/Å^2$) | 45.156 | |
| Pixel size (Å) | 0.825 | |
| Number of micrographs | 9,352 | |
| Final particle number | 8,145 | 3,117 |
| Symmetry imposed | I | C1 |
| Map resolution (Å) (FSC=0.143) | 2.21 | 19.35 |
| Map resolution (Å) (FSC=0.5) | ND | ~21 |
| Map resolution range (Å) | 1.82-35 | NA |
| Map sharpening B-factor ($Å^2$) | 61.0 | |
| **Model** | | |
| PDB | | |
| Model Composition<br>Chains | <br>3 | |

| | |
|---|---|
| Non-hydrogen atoms<br>Protein residues<br>Ligands | 7,681<br>1,028<br>0 |
| R.m.s. deviations<br>Bond lengths (Å)<br>Bond angles (°) | <br>0.003<br>0.546 |
| Validation<br>MolProbity score<br>Clashscore<br>Poor rotamers (%)<br>Cβ outliers (%)<br>CaBLAM outliers (%)<br>Cis proline / general (%)<br>Twisted proline / general (%) | <br>0.84<br>1.22<br>0.00<br>NA<br>1.48<br>8.3/0.0<br>0.0/0.0 |
| Rama-Z<br>Whole (N = 1022)<br>Helix (N = 136)<br>Sheet (N = 298)<br>Loop (N = 588) | <br>0.43 (0.27)<br>0.57 (0.44)<br>0.35 (0.30)<br>0.32 (0.27) |
| Ramachandran plot (%)<br>Favored (%)<br>Allowed (%)<br>Outliers (%) | <br>98.24<br>1.76<br>0.00 |
| B-factors (Å$^2$) (min/max/mean) | 17.83/82.66/34.59 |
| **Data** | |
| d99 masked (full/half1/half2) | 2.35/1.69/1.69 |
| d99 unmasked (full/half1/half2) | 2.28/1.67/1.67 |
| FSC (model) = 0 (masked/unmasked) | 2.11/2.18 |
| FSC (model) = 0.143 (masked/unmasked) | 2.18/2.21 |

| | |
|---|---|
| FSC (model) = 0.5 (masked/unmasked) | 2.31/2.79 |
| **Model vs Data** | |
| CC (mask) | 0.91 |
| CC (box) | 0.47 |
| CC (peaks) | 0.33 |
| CC (volume) | 0.89 |
| EMRinger score | 6.43 |
